# Metapopulation connectivity in Voles (*Microtus* sp.) as a gauge for tallgrass prairie restoration in midwestern North America

**DOI:** 10.1101/2020.10.17.343301

**Authors:** Marlis R. Douglas, Whitney J.B. Anthonysamy, Mark A. Davis, Matthew P. Mulligan, Robert L. Schooley, Wade Louis, Michael E. Douglas

## Abstract

Applying quantifiable metrics to validate the success of restoration efforts is crucial for ongoing management programs in anthropogenically fragmented habitats. Estimates of dispersal can provide such baseline data because they measure not only the extent to which restored patches are colonized and interconnected, but also their metapopulation source/sink dynamics. In this context, we estimated dispersal and population connectivity among prairie (*Microtus ochrogaster*; N=231) and meadow vole (*M. pennsylvanicus*; N=83), sampled from eight restored plots at five tallgrass prairie sites embedded within the agricultural matrix of midwestern North America. Our expectation was that extensive distances separating these restored habitats (i.e., 48–246 km) would spatially isolate vole metapopulations, resulting in significant genetic differentiation. We first used molecular taxonomy to validate the field-identifications of all sampled individuals, then used pairwise *F_ST_* derived from 15 microsatellite DNA loci to estimate genetic connectivity among the species-delimited study populations. Metapopulation stability was gauged by assessing migration rates and deriving effective population sizes (*Ne*). We also calculated relatedness values (*r*) as a potential surrogate for contact in prairie vole, a primary vector for Lyme disease. Molecular species-assignments contravened field-identifications in 25% of samples (11 prairie/67 meadow) and identified two instances of species-hybridization (0.6%). Local effects (i.e., population crash/drought) were manifested at two sites, as documented by significant temporal declines in *Ne* and *r*. Overall, high migration rates and non-significant (10/15) pairwise *F_ST_* values underscored elevated metapopulation connectivity. A single site that recorded five significant *F_ST_* values also displayed significant *r*-values indicating the inadvertent sampling of closely related individuals. This highlights the close social groupings among cooperatively-breeding prairie vole that can exacerbate Lyme disease transmission. Thus, while elevated population connectivity aligns with prairie restoration goals, it also reinforces a need in adaptive management to evaluate environmental matrices for their permeability to vector-borne diseases.

## Introduction

‘Distribution’ and ‘abundance’ are not only recognized as natural history attributes of species, but also the building blocks of ecology [1]. Yet, their functional integrity is being seriously eroded by an anthropogenic fragmentation of habitat [2], with the rendering of species distributions into subdivided, often isolated populations as a result of large-scale modifications. Ancillary considerations, such as ‘dispersal’ and ‘population connectivity,’ have instead become paramount [3], in that they more appropriately mirror the effects of the Anthropocene [4]. These impacts are most apparent within temperate grassland biomes (>8% of the global landmass [5]) reflecting the ease with which these biomes can be transitioned from their native state into anthropogenic settlements and agricultural plots.

The anthropogenic conversion of grasslands is most apparent in the Tall Grass Prairie of Midwestern North American [6], where greenhouse gas emissions exceed the national average by >20 [7]. Furthermore, the growing season and agricultural row-crop technology inherent to this region have now been significantly extended ([8], table 1), and the anthropogenic magnification both components has subsequently enhanced the already well-established agricultural capacity of the region [9]. In addition, an increasing market demand for biofuels has intensified the removal of perennially rooted vegetation, leading to the general elimination of potential ‘edge’ habitats along the margins of agricultural fields that served to sustain essential connectivity among habitat fragments [10]. Other consequences of large-scale habitat alterations are the loss of indigenous biodiversity and biotic homogenization [11].

**Table 1.** Total number of vole samples collected during three field seasons (2010–2012) across five Illinois SAFE sites. Samples were harvested non-invasively from two species [prairie vole (*Microtus ochrogaster*) and meadow vole (*M. pennsylvanicus*)], but only totals are listed. Figure 1 shows geographic location of sites.

| Site | 2010 | 2011 | 2012 | Totals |
| --- | --- | --- | --- | --- |
| Livingston | 27 | 4 | 10 | 41 |
| Montgomery | 0 | 56 | 41 | 97 |
| Pontiac | 21 | 1 | 9 | 31 |
| Prairie Ridge | 60 | 3 | 32 | 95 |
| Saybrook | 0 | 19 | 77 | 96 |
| Total | 108 | 83 | 169 | 360 |

The detrimental consequences of these anthropogenic impacts can be stemmed, or even reversed through habitat restoration. In the prairie landscape of North America, numerous conservation initiatives have been implemented so as to improve remnant grassland parcels and increase population numbers of targeted wildlife. This is manifested in Illinois through project SAFE (State Acres for Wildlife Enhancement; http://www.dnr.state.il.us/orc/safe/), a managerial approach actively promoted by the Illinois Department of Natural Resources (IDNR) as a means of rehabilitating agricultural land specifically within two targeted regions: Grand Prairie Natural Division (east-central IL), and Southern Till Plain (south-central IL).

However, one critical aspect of restoration is how ‘success’ is defined, and what metrics are potentially used to gauge its success. As a mechanism to quantify the effectiveness of restoration efforts, we gauged impacts of habitat rehabilitation on metapopulation connectivity of prairie vole (*Microtus ochrogaster*) and meadow vole (*M. pennsylvanicus*), as a means of evaluating the spatio-temporal genetic structure of this particular biodiversity component. Herein, metapopulations are defined as a collection of spatially separated subpopulations that experience some level of gene flow [12]. Additionally, voles are a model organism to evaluate the manner by which habitat quality can impact population dynamics and demographics [13]. They are easily captured via live trapping, a technique that yields replicable population size estimates. Voles can also subsist in smaller home ranges (100m^2^) within relatively reduced habitats, yet still reflect population-scale processes. Grasses and sedges are preferred habitats, and this is beneficial from an experimental stance in that the habitat can be easily manipulated so as to produce patches of different quality and quantity, thus promoting hypothesis testing within a field setting [14]. Reduced ground cover, for example, tends to impede dispersal, reduce residency, and promote mortality due to predation. In addition, landscapes artificially modified to accommodate a diversity of habitat-types also allow the imposition of experimental designs that are difficult to establish and/or maintain in a natural setting. Live trapping associated with fenced plots, for example, can separate dispersal from mortality, a consideration difficult to obtain under natural conditions. It also allows for iterative replication, again difficult to accomplish when study plots are open landscapes.

We evaluated vole metapopulations across five widely distributed SAFE sites deeply imbedded within an anthropogenic row-crop landscape as a mechanism to test if large topographical distances, coupled with the extent to which patches are separated within the highly modified agricultural matrix, would act synergistically to curtail vole dispersal [15]. Because prairie vole is a highly philopatric and socially monogamous rodent that often breeds cooperatively [16], it offered an opportunity to explore the manner by which social- and kin-structure in vole populations might impact dispersal.

We employed microsatellite DNA analyses to 1) quantify genetic diversity in both vole species within and among the SAFE sites, and 2) assess levels of temporal and spatial gene flow as a metric for landscape permeability. In this sense, population genetic patterns reflect dispersal of organisms through complex environments, and hence can reveal if connectivity indeed exists among patches widely separated within an anthropogenically-modified landscape [17]. This, in turn, provides an estimate for the permeability of the landscape, a relevant aspect in that diverse patches interspersed with edges and corridors are important components for conservation and management within agro-ecosystems [18]. We expected metapopulation structure to be defined by limited connectivity among study sites, particularly given the life histories of our study species and the dominance of the agro-ecosystem within which our study plots were isolated.

## Materials and methods

### Sampling and DNA extraction

Prairie and meadow vole were live trapped from mid-May through mid-August (2010–2012) across five Illinois SAFE sites (Fig 1). Sampling plots within each (i.e., 1.96–64.58ha) contained six transects of 15 traps set 7m apart, with trapping conducted over three consecutive evenings (N=90 traps/ plot/ night [19]). Ear tissue was sampled from each captured vole and stored for genetic analyses in 95% EtOH at −20°C. Whole genomic DNA was extracted using Promega Wizard Kit (2010–2011) or Qiagen DNeasy Blood and Tissue Kit (2012) and quantified using an Implen Pearl P-300 nanophotometer.

**Fig 1.**
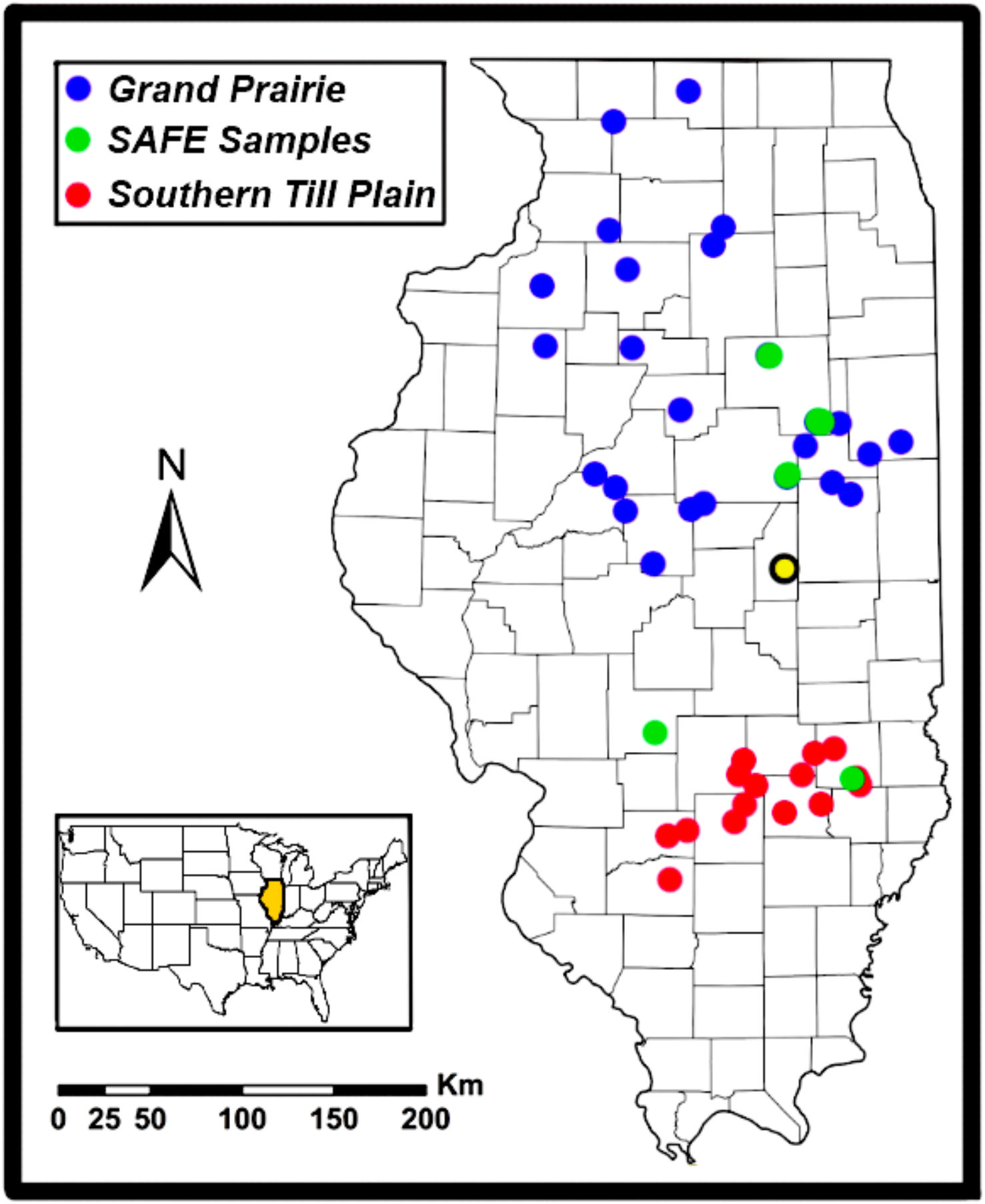
Map depicting rehabilitation locations for Illinois State Acres for Wildlife Enhancement. (SAFE: http://www.dnr.state.il.us/orc/safe/). Blue indicates SAFE sites located in the Grand Prairie, and red sites in the Southern Till Plain. SAFE sites sampled for prairie vole (*Microtus ochrogaster*) and meadow vole (*M. pennsylvanicus*) during 2010–2012 are depicted in green. Yellow indicates study location documenting infestation of *M. ochrogaster* by blacklegged or deer tick (*Ixodes scapularis*) and its Lyme disease bacterium, *Borrelia burgdorferi* (Bb).

### Microsatellite DNA Genotyping

Microsatellite loci previously developed for vole species were evaluated for consistent cross-amplification in prairie and meadow vole, with as set of 23 loci selected and combined into six multiplex panels using fluorescently labeled forward primers (Supplemental Appendix I). Amplifications were conducted in 10–15μl volume polymerase chain reactions (PCR) using approximately 10–15ng template and following standard procedures.

#### Fragment Analysis

Microsatellite DNA fragments were resolved on an automated Applied Biosystems (ABI) Prism 3730xl GeneAnalyzer (W. M. Keck Center, University of Illinois, Champaign). An internal size standard (Liz 500) was run with each sample, and alleles were scored using GeneMapper© 4.1 software (ABI). Genotypes were partitioned by species, site, and sampling period, then tested via Micro-Checker 2.2.3 [20] for possible null alleles, large allele dropout, and scoring errors. All pairs of loci were tested for linkage disequilibrium (Markov Chain parameters: 10,000 dememorization steps, 500 batches, 5,000 iterations), and each locus evaluated for deviations from Hardy-Weinberg equilibrium (HWE) using exact tests (GenePop 4.0; [21]) with Bonferroni correction for multiple comparisons [22].

#### Assignment of individuals to species

Prairie and meadow vole are morphologically similar, and accurate identification can be difficult particularly when non-lethal sampling is conducted. To verify field-based morphological species identification, we first used the population assignment option in GenAlex 6.50 [23] with samples clustered according to similarities in genotype. Genetic-based species assignments were used to re-classify individuals to species that were then re-evaluated for distinct gene pools using a Bayesian assignment test (Structure 2.3.4 [24]), with admixture ancestry and correlated allele frequency options employed [25].

#### Molecular taxonomy

To further verify species designation, a diagnostic locus (*Microtus avpr1a* gene [26]) was evaluated that displays considerable size differences between the two species, with prairie vole at ~600–800bp and meadow vole at ~200–300bp. A subset of samples (n=68) was tested for concordance between field- and genetic-based species identification, using primers and PCR protocols adapted from previous studies [27,28,29]. Species identification was confirmed by separating PCR products on 2.5% agarose gel, staining with GelGreen (Biotium Inc., Hayward, CA), and visualization on a bluelight transluminator.

### Genetic diversity and structure of vole metapopulations

Standard population genetic parameters were calculated as quantitative metrics for diversity within and among sites. Allelic frequencies, observed (*H*_O_), and expected heterozygosities (*H*_E_) were estimated for each species at each site using GenAlEx. Values for allelic richness (A_R_) and private allelic richness (P_AR_) were derived from rarefaction and corrected for sample sizes (HP-Rare v. June-6-2006 [30]). In addition, pairwise relatedness (r) was evaluated among individuals using the Ritland-1996 estimator in GenAlex, so as to reduce potential bias in genetic estimates caused by inadvertent sampling of related individuals in localized trapping transects.

#### Bayesian clustering

To further assess distribution of gene pools among and within sampling locations, we conducted assignment tests using Structure. Ten replicates were run for *K*-values ranging from one to the number of sampling locations or years, using a burn-in of 500,000 iterations followed by 1,000,000 Markov Chain Monte Carlo (MCMC) replicates. The optimal number of clusters for each simulation (per Structure Harvester 0.6.1 [31]) then allowed a calculation of the *ad hoc* statistic ‘ΔK’ [32]. To test for isolation by distance (IBD), a Mantel test was conducted in GenAlEx using 999 permutations. Genetic divergence was gauged as *F_ST_*/(1-*F_ST_*), and geographic distances (log-transformed) were evaluated as the shortest distance between sites (ArcMap 10.1; ESRI 2012, Redlands, CA, USA).

#### Pairwise *F_ST_*

We employed *F_ST_*-values as a standardized index to gauge population connectivity and as indicators of gene flow, with values varying from 0 (identical allele frequencies indicating panmixia) to 1 (populations fixed for different alleles indicating isolation) [12]. Spatial and temporal patterns of gene flow were estimated by plot (transect) among sites within years, and within sites among years (GenAlEx, with 9,999 permutations and missing data interpolated). Gene flow could only be reliably assessed for prairie vole, due to variability in numbers of samples obtained at sites among years, as well as the uneven distribution of samples following genetic re-classification to species.

Pairwise *F_ST_* values for prairie vole were derived among three SAFE sites (i.e., Livingston, Pontiac, and Prairie Ridge) in 2010, and two sites in 2012 (i.e., Montgomery and Prairie Ridge). Pairwise *F_ST_* was similarly derived among years across two sites subjected to environmental perturbations (Montgomery: 2011 *versus* 2012/ Prairie Ridge: 2010 *versus* 2012).

### Estimates of migration and effective population size

Individual movements heavily influence metapopulation dynamics, and two are of particular importance in this regard: Colonization (i.e., movement into a novel habitat patch) and migration (i.e., the immigration into occupied patches). For example, the annual replanting of crops is an agro-ecosystem dynamic that annually converts vole subpopulations into sinks that must be potentially re-colonized from a source [33].

#### Recent migration among SAFE sites

Potential F1 descendants of migrant individuals were identified using GeneClass2 2.0 [34]. We selected the ‘L_home’ test statistic (i.e., the likelihood of obtaining the genotype of an individual from the sampled population), under the assumption that not all source populations were sampled. A Monte Carlo resampling method was selected, with 10,000 bootstraps and a threshold value of 0.05, as it offers improved performance when identifying first generation migrants while also controlling for Type-I error rates [35].

#### Effective population size

Trapping success was uneven among years, impacted by an apparent population crash in 2011 and a substantial drought in 2012 [19]. To gauge the effects of these perturbations on vole metapopulations, we quantified effective population size (*Ne*) [36] based upon estimates of linkage disequilibrium (i.e., a non-random association of independent alleles with haplotypes occurring in unexpected frequencies [37]). The *Ne* metric reflects the rate of random genetic drift (i.e., random fluctuations in allele frequencies over time [12]), as well as the effectiveness of selection and migration. It also indicates the loss of heterozygosity in the population, and links strongly with demographic factors such as sex ratio, population size, and lifetime fitness.

Due to sample size limitations, *Ne* could only be derived for prairie vole at Prairie Ridge (2010 *versus* 2012) and Montgomery (2011 *versus* 2012). Mean *Ne* was compared by year at these two SAFE sites by implementing Welch’s t-test for unequal variances. This approach is more robust than Student’s t-test and maintains Type-I error rates despite inequalities among variances and sample sizes. The test was performed in R [38] using summary statistics and a suitably modified web-based program (http://stats.stackexchange.com/questions/30394/how-to-perform-two-sample-t-tests-in-r-by-inputting-sample-statistics-rather-tha).

## Results

### Sampling, genotyping, and species assignments

Over the three field seasons, we acquired a total of 360 samples across five sites (Table 1). Of these, 194 were field-identified as prairie vole, and 166 as meadow vole, yielding a prairie-to-meadow-vole ratio of 1.17/ 1.0. All samples were genotyped across 23 microsatellite loci, with eight loci subsequently removed, due either to null alleles or scoring problems, with 15 loci remaining for data analyses. In addition, 46 samples were excluded due to the presence of missing data, leaving N = 314 for detailed evaluations. Linkage disequilibrium was detected but was inconsistent across temporal periods or sites, and thus attributed to demographic effects on genetic structure rather than non-independence amongst loci.

Genotype-based species assignment was concordant with 75% of field identifications based on morphology (236 of 314). Among the remaining 78 field identifications, 11 prairie and 67 meadow voles were genetically reclassified (Fig 2, top), resulting in a 231 prairie and 83 meadow voles, respectively. A subsequent Bayesian assignment test based on genetic species identification consistently allocated all 314 samples (Fig 2, bottom). Screening with the diagnostic *avpr1a* locus confirmed species-level classification for 97% of test samples (65/67), with two individuals (0.6% of 314) identified as admixed (Bayesian assignment plot; Fig 2, bottom), suggesting in turn a rare occurrence of hybridization between species. Genetic species assignments rather than field identification were employed in subsequent analyses.

**Fig. 2.**
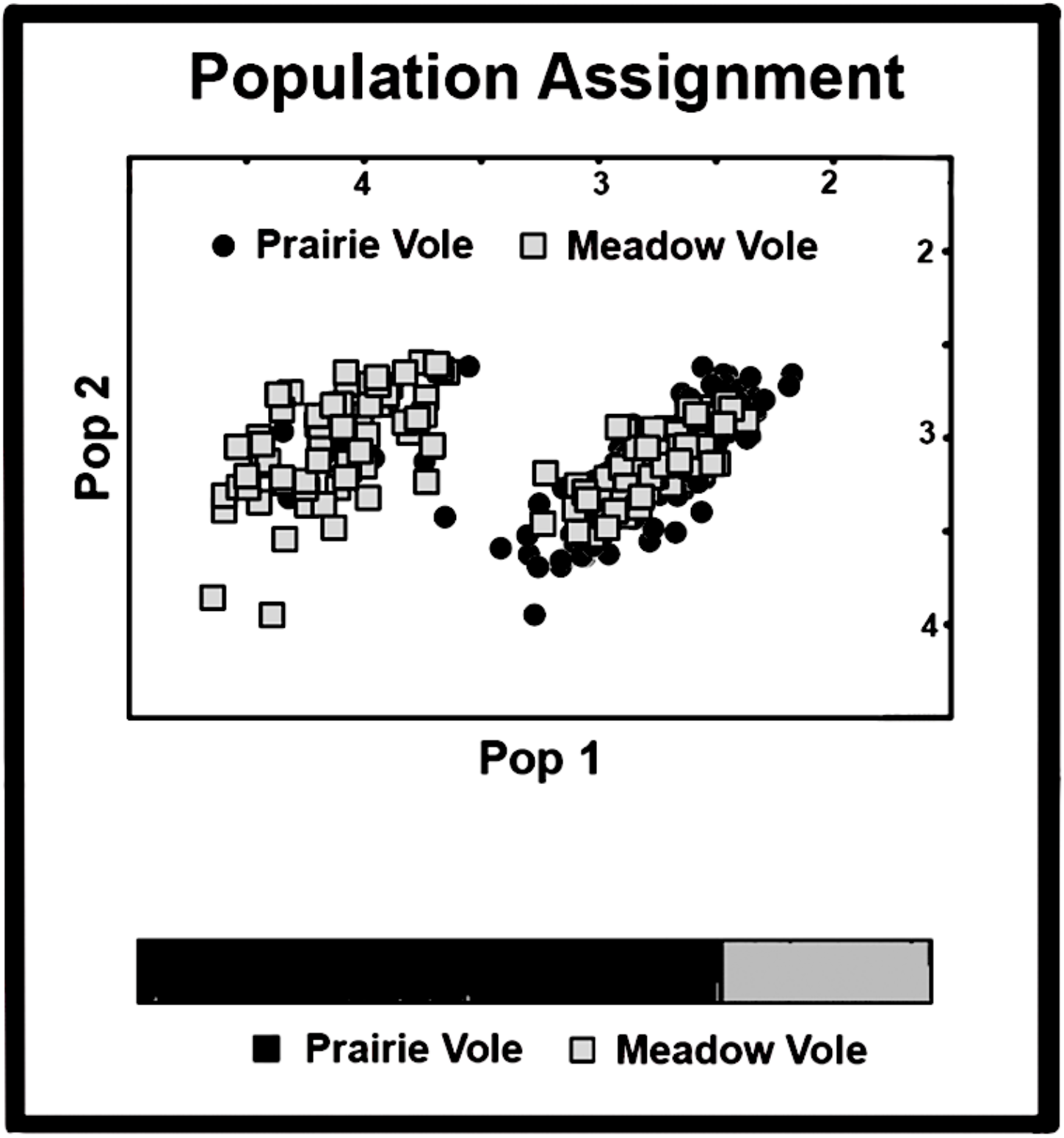
Top: Samples of prairie vole (*Microtus ochrogaster*) and meadow vole (*M. pennsylvanicus*) allocation to species. Samples (N=314) were collected at five Illinois SAFE sites from 2010–2012. Initial species identification in the field was based on morphology. Genotypes, derived across 15 microsatellite loci, were then used to allocated samples to two gene pools using a population assignment test in GenAlEx v. 6.5. Bottom: Allocation of genotypes derived from 15 microsatellite loci using a population assignment test in Structure v2.3.4. Species identification was based on molecular genetic reclassification. Shading reflect distinct gene pools and vertical bars represent probabilities of assigning an individual to a gene pool.

### Genetic diversity and metapopulation structure

Microsatellite DNA polymorphism was high in both species, with an average of 22.7 and 16.3 alleles per locus, with observed heterozygosity (*H*_O_ for prairie and meadow vole) = 0.80 and 0.70, respectively (Table 2). Pairwise *F_ST_* values were non-significant for prairie vole by plot and site, at both spatial and temporal scales, save for comparisons with Prairie Ridge (Table 3). These significant *F_ST_* values were not associated with geographic distance, and a test for IBD was non-significant (*p*=0.36).

**Table 2.**
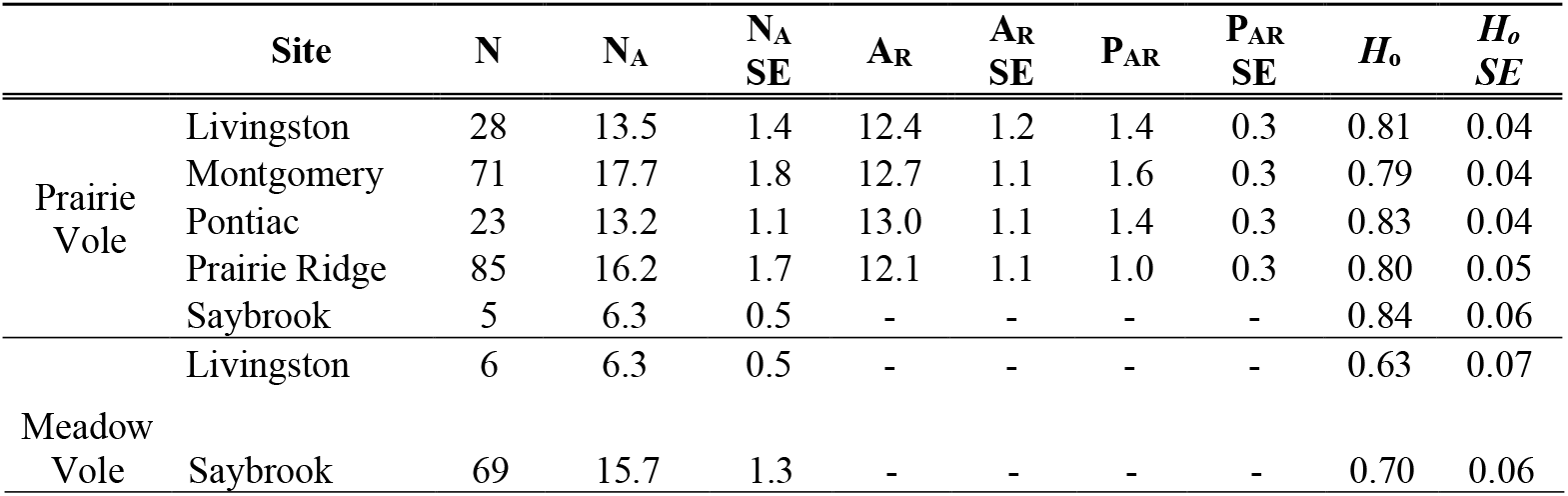
Diversity statistics based on 15 microsatellite loci for 212 prairie vole (*Microtus ochrogaster*) and 75 meadow vole (*M. pennsylvanicus*) collected from five Illinois SAFE sites. Listed are: N = number of individuals genotyped per locus; N_a_ = average number of alleles per locus; Na SE = Standard error for average number of alleles per locus; A_R_ = allelic richness corrected for sample size; A_R_ SE = Standard error for allelic richness corrected for sample size; P_AR_ = private allelic richness corrected for sample size; P_AR_ SE = Standard error for private allelic richness corrected for sample size; *H*_O_ = observed heterozygosity; *H*_O_ SE = Standard Error for observed heterozygosity.

**Table 3.** Pairwise *F_ST_* estimates derived from 15 microsatellite DNA loci for prairie vole (*Microtus ochrogaster*) collected by year from plots (=Plot) in Illinois SAFE sites (=Site). *F_ST_* values are below diagonal and P-values above diagonal. Bonferroni-adjusted statistical significance for 2010 (upper panel) = p<0.003, and P<0.008 for 2012 (lower panel). Sample sizes for plots are in Table 5. Significant values are in bold.

| 2010 - Site | Plot | L-H | L-M | P-C | P-T | PR-H | PR-T |
| --- | --- | --- | --- | --- | --- | --- | --- |
| Livingston | Hummel | X | 0.893 | 0.024 | 0.475 | 0.079 | **0.002** |
| Livingston | Marge | 0.020 | X | 0.601 | 0.783 | 0.580 | **0.001** |
| Pontiac | Curve | 0.031 | 0.028 | X | 0.923 | 0.156 | **0.001** |
| Pontiac | Tower | 0.037 | 0.036 | 0.036 | X | 0.534 | 0.090 |
| Prairie Ridge | Harvey | 0.017 | 0.017 | 0.022 | 0.029 | X | **0.001** |
| Prairie Ridge | Tombstone | 0.025 | 0.029 | 0.033 | 0.038 | 0.018 | X |

| 2012 - Site | Plot | M-H | M-L | PR-H | PR-T |
| --- | --- | --- | --- | --- | --- |
| Montgomery | Huber | X | 0.023 | **0.006** | 0.012 |
| Montgomery | Lane | 0.025 | X | 0.011 | **0.001** |
| Prairie Ridge | Harvey | 0.055 | 0.044 | X | 0.013 |
| Prairie Ridge | Tombstone | 0.027 | 0.023 | 0.043 | X |

Assignment tests revealed three distinct gene pools (Fig 3) that were primarily distributed across sites, with spatial substructure evident within but one site (i.e., Prairie Ridge; Fig 3A and Fig 3B). Interestingly, substructure at this site was retained from 2010 to 2012 (Fig 3C), but ancestry proportions shifted significantly between years following the 2011 population crash (*F_ST_* = 0.014; *P*<0.0001), a pattern consistent with either drift or re-colonization. In contrast, the second site (i.e., Montgomery; Fig 3D) showed almost identical ancestry proportions in gene pools for 2011 and 2012.

**Fig 3.**
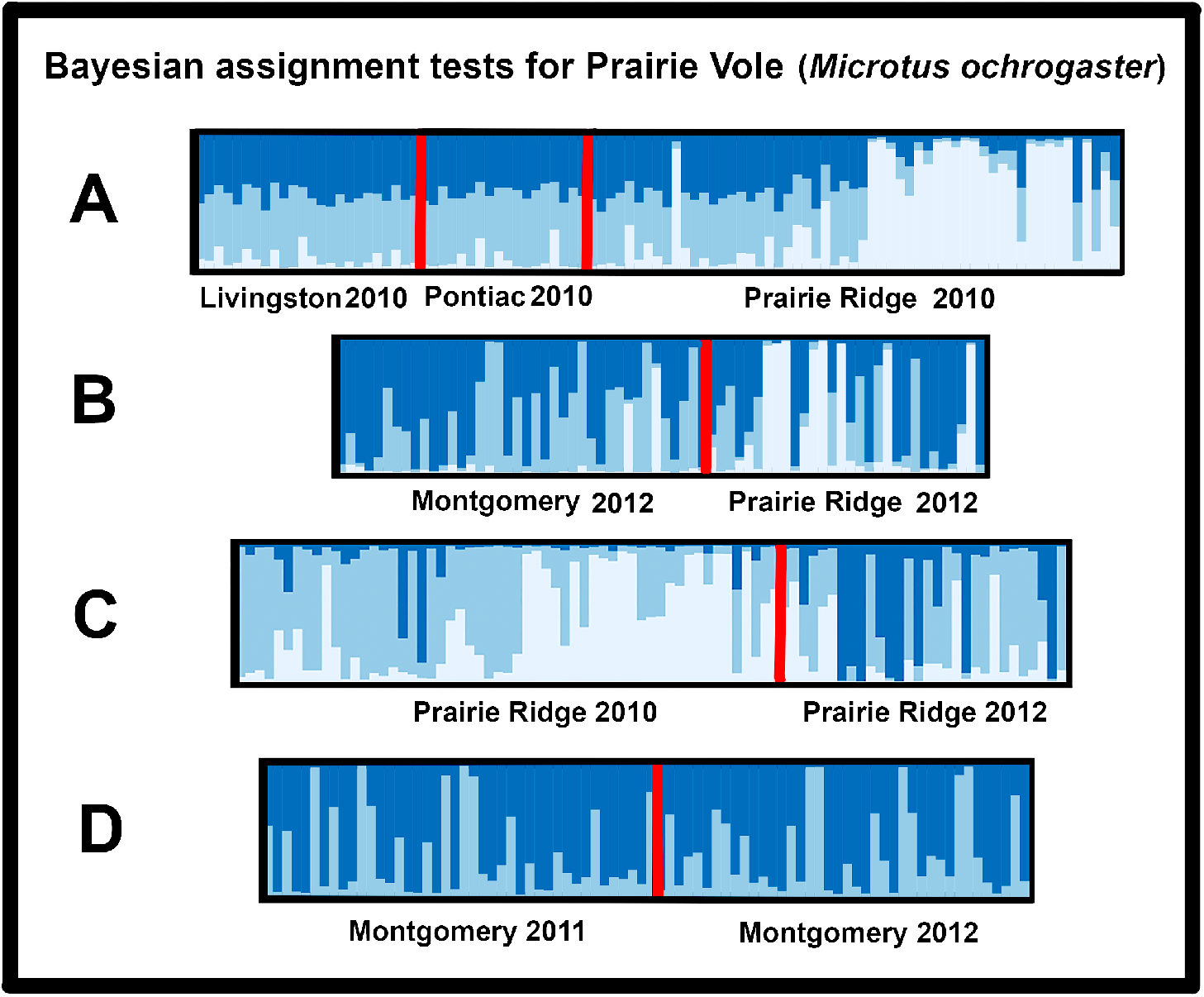
Spatial patterning of gene pools within Prairie Vole (*Microtus ochrogaster*) among Illinois SAFE sites in A) 2010 and B) 2012, and among years within Illinois SAFE sites C) Prairie Ridge and D) Montgomery. Plots are based on genotypes derived across 15 microsatellite and based on Bayesian Assignment tests (Structure v2.3.4). Colors reflect distinct gene pools and vertical bars representing individuals and proportion of colors reflecting probabilities of ancestry in a gene pool. Sample sizes are in Table 5.

### Estimates of migration and effective population size

Estimates of migration calculated in GeneClass2 indicated contemporary movements of individuals among SAFE sites (Table 4). In prairie vole, five (2%) individuals had a probability threshold below 0.01 and were identified as potential migrants based on assignment of one sample from Livingston to Pontiac, two from Montgomery to Livingston, one from Pontiac to Livingston, and one individual from Prairie Ridge to Pontiac. No potential migrants were identified for meadow vole, but this species was only captured at two sites, with six individuals sampled from Livingston and the remainder from Saybrook.

**Table 4.**
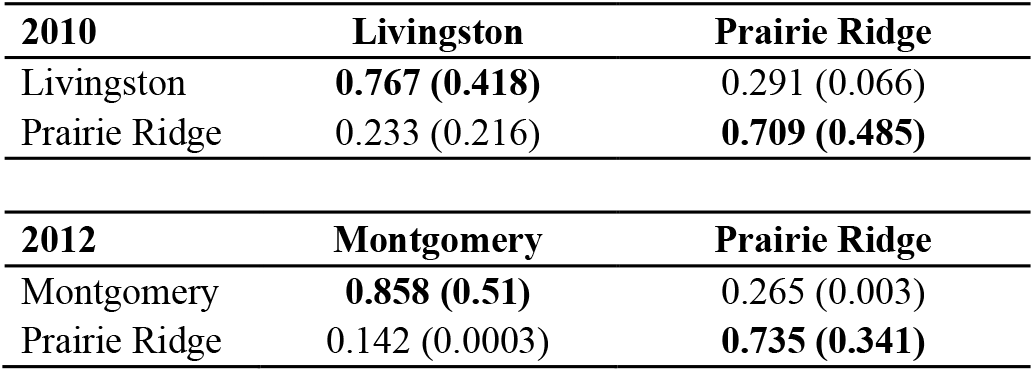
Estimated rates of migration *versus* residency calculated for prairie vole (*Microtus orchrogaster*) within and between three Illinois SAFE sites (Livingston, Montgomery, and Prairie Ridge). Vole samples were collected over two years: 2010 [Livingston (N=24); Prairie Ridge (N=57)] and 2012 [Montgomery (N=35); Prairie Ridge (N=30)]. Values are posterior estimates for mean and 95% highest posterior density interval (in parentheses). Bold values = residency estimates.

#### Population crashes

In 2011, effective population size (*Ne*) in prairie vole declined significantly from 119 at Prairie Ridge in 2010 (95% CI = 84-195) to 36 in 2012 (95% CI = 28-50) (t = 4.58; P<0.00002). Prairie vole displayed significantly lower *Ne* estimates during the 2012 drought as well, with Montgomery declining from 1,314 (95% CI = 180-∞) in 2011 to 119 (95% CI = 80-221) in 2012 (t = 3.06; P<0.0032).

### Relatedness

Average relatedness (*r*) within plots ranged from 0.009–0.106, but values were generally higher among individuals at Prairie Ridge than at other sites (Table 5). Relatedness at the Harvey plot differed significantly from 2010 to 2012 (t=-3.06, *P*<0.028), whereas those for the Tombstone plot did not (t=-0.54, *P*>0.6). Across plots and sites, the average pairwise relatedness and geographic distance were significantly but inversely related in 2010, with relatedness diminishing as distances increased (3 sites, 6 plots; *P*=0.027). However, relatedness *versus* distance was non-significant in 2012 (2 sites, 4 plots; *P*=0.13).

**Table 5.**
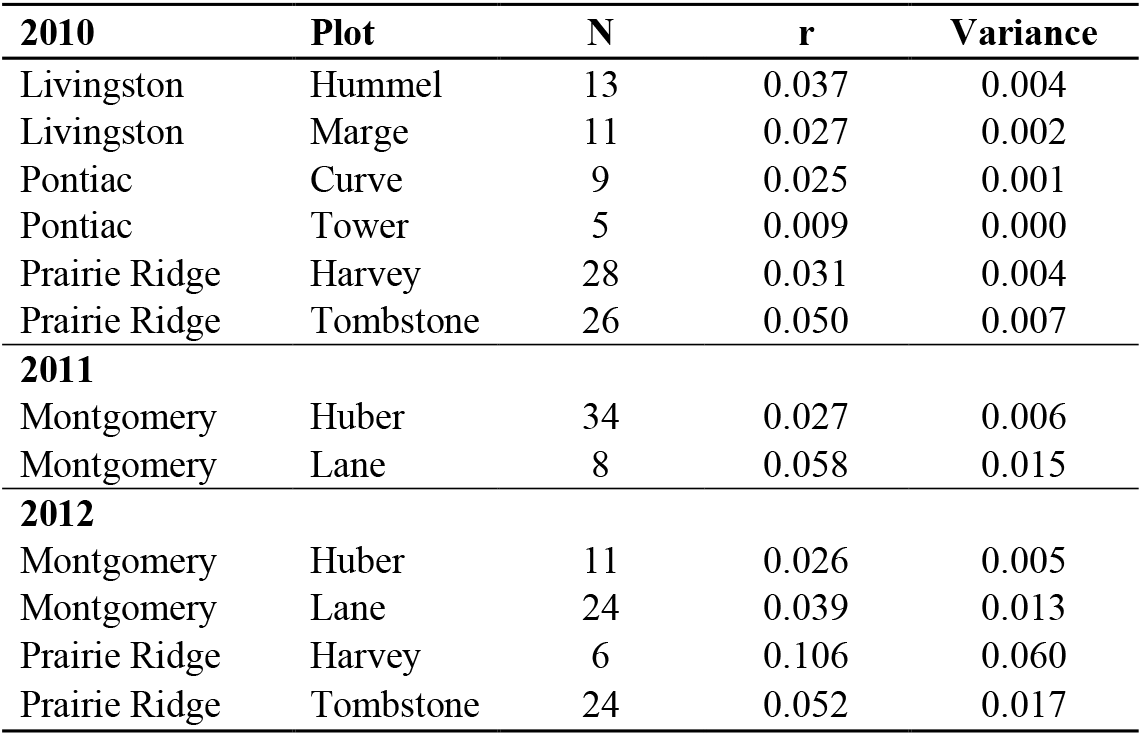
Yearly estimates of average relatedness (r) and variance derived from 15 microsatellite DNA loci from samples (=N) of prairie vole (*Microtus ochrogaster*) collected by plot (=Plot) in Illinois SAFE sites (= Livingston, Pontiac, Montgomery, and Prairie Ridge)

| **2010** | **Plot** | **N** | **r** | **Variance** |
| --- | --- | --- | --- | --- |
| Livingston | Hummel | 13 | 0.037 | 0.004 |
| Livingston | Marge | 11 | 0.027 | 0.002 |
| Pontiac | Curve | 9 | 0.025 | 0.001 |
| Pontiac | Tower | 5 | 0.009 | 0.000 |
| Prairie Ridge | Harvey | 28 | 0.031 | 0.004 |
| Prairie Ridge | Tombstone | 26 | 0.050 | 0.007 |
| **2011** |  |  |  |  |
| Montgomery | Huber | 34 | 0.027 | 0.006 |
| Montgomery | Lane | 8 | 0.058 | 0.015 |
| **2012** |  |  |  |  |
| Montgomery | Huber | 11 | 0.026 | 0.005 |
| Montgomery | Lane | 24 | 0.039 | 0.013 |
| Prairie Ridge | Harvey | 6 | 0.106 | 0.060 |
| Prairie Ridge | Tombstone | 24 | 0.052 | 0.017 |

## Discussion

An essential consideration in the management of anthropogenically fragmented habitats is to gauge the success of ongoing restoration efforts. Metrics are needed to quantify the extent to which restored patches are colonized and interconnected. The metapopulation dynamics of small mammals are often used to gauge habitat fragmentation [39], and conversely, the potential benefits accrued from anthropogenic land conversion [40]. However, large, homogenized agricultural landscapes are less well studied in this regard, due largely to a lack of community diversity when compared to natural habitats [41]. In addition, those studies that previously evaluated genetic structuring and diversity in agro-ecosystems reported conflicting results. For example, values can often exceed those found in more uniform natural areas, particularly given the aforementioned variability induced by agricultural management [42]. Conversely, population dynamics of voles can supersede the annual perturbations found in an agricultural landscape, such that isolation-by-distance (IBD) becomes the only delimiter for gene flow [43].

Our empirical evaluation of vole population genetic structure provides additional insights into these issues, and particularly those mechanisms by which restored patches of an agro-ecosystem are interconnected. Our results have important management considerations that impinge directly upon the achievement of habitat restoration goals. They may also have subsidiary ecological implications, particularly with regard to the permeability of an environmental matrix to vector-borne diseases [44].

### Taxonomic uncertainty and species abundances

The correct identification of species in the field is a fundamental assumption when ecological hypotheses are tested, and when conservation and management initiatives are implemented. Thus, by ignoring species uncertainty particularly among *Microtus* spp., biased estimates regarding population processes can emerge [45]. Meadow and prairie vole can be difficult to distinguish in the field, and their differentiation is based on qualitative characters that relate to pelage, as well as differences in tooth morphology [46].

Fortunately, species-diagnostic DNA markers can clearly discriminate among sympatric species [47], as well as decipher potential hybridization and introgression. For example, we found a 25% mismatch between field-identifications and those derived genetically for prairie and meadow vole, and similar disparities have also been observed in other studies [26]. We also documented two samples at one site (i.e., Saybrook) that represented hybridized or introgressed individuals. Meadow vole experienced a significant range expansion from northern into central Illinois during the 1970’s, facilitated by the construction of interstate highways and accompanying right-of-ways. The impacts on Illinois wildlife have varied [48]. In this instance, it created a local contact zone between the two vole species that impacted their distributions [49]. These aspects underscore the practicality of molecular species identification, particularly in our example with regard to conflicting identification among sympatric voles.

Prairie and meadow vole respond differently to the composition of the environment, particularly as it relates to trophic and habitat preferences [50,51]. Prairie vole is more tolerant of sparse cover, for example, whereas meadow vole prefers more dense vegetation. Habitat preference may explain the disparity in our samples with regard to their presence and/ or abundance, particularly given that vegetation communities varied among study sites [19]. Because voles are important grassland components [52] with responses that differ with regard to vegetation management, their accurate identification is clearly an important consideration when evaluating the success of prairie restoration. Our study demonstrates the effectiveness of genetic data in accurately diagnosing species.

### Dispersal capacity in agricultural landscapes

Small mammal communities are substantial components of grassland biodiversity and are intimately linked with complex trophic interactions in grassland ecosystems [52]. Deer mice (*Peromyscus* sp.) and vole (*Microtus* sp.) are ubiquitous in these systems and their metapopulation dynamics are strongly influenced by crop type, farming practices, and vegetational structure [51].

Most habitats are characterized by spatial heterogeneity with regard to patch quality and this variability modulates demographic processes, such as population density and dispersal rates. Experimental manipulations of patch quality (with regard to amount of food and predation risk) demonstrate that patch quality affects dispersal in prairie vole [53], as well as their intrinsic population demographics (e.g., growth rates and reproductive success; [54], but not social structure [55]).

The impacts of habitat fragmentation on movements of small mammals were examined in a 7-year mark-recapture study conducted over three types of 0.5ha blocks: Single large patches (5,000m^2^); clusters of medium patches (288m^2^); and clusters of small patches (32m^2^) [14]. Prairie vole moved further as fragmentation increased, but did so in decreasing proportions, with male dispersal rates exceeding those of females. Also, a greater number (primarily juvenile males) switched from smaller blocks in the study design to larger blocks. The researchers concluded that, at the small scale of 5,000m^2^, most species in fragmented habitats moved farther when they did move (presumably for trophic and reproductive purposes), but did not move often, potentially due to costs associated with distances traveled.

We predicted the highly fragmented and agriculturally dominated landscape of central Illinois would impede vole dispersal, particularly given that SAFE sites were separated by 48–246km from one another. Yet, prairie vole displayed a greater than expected capacity for dispersal, as evidenced by both temporal and spatial population structure, suggesting in turn that gene flow is clearly, and surprisingly, maintained across study sites. ‘Border habitats’ such as fencerows, roadside ditches, and waterways can promote re-colonization of restored grasslands [56], and in this regard, may be especially important in maintaining vole connectivity, particularly given their rapid demographic responses that often follows population crashes [57]. In this sense, restored grasslands can serve as a ‘source’ in the context of vole metapopulation dynamics, and thus buffer those subsequent population declines that are induced by the annual harvesting/ replanting of crops [19].

### Genetic structure in voles

Gene flow in prairie vole is also modulated spatially and temporally, as observed in other studies that examined spatial and scale-dependent variability in gene flow [33]. *F_ST_* and Structure analyses showed weak divergence among sites, and admixture within sites was greater for consecutive (i.e., 2011 *vs*. 2012) than non-consecutive years (i.e., 2010 *vs*. 2012).

Geographic distance can be a primary factor in determining the dispersal of rodents in converted ecosystems [58]. However, the relationship between geographic distance and patterns of genetic structure was non-significant in our study. Although SAFE sites demonstrated high connectivity, as evidenced by a lack of strong population structure and the potential for migrant individuals among sites, the subtle structure detected by *F_ST_* analyses could indicate a relationship between large population size and long-distance dispersal as imposed by the agroecosystem matrix [43]. Further, temporal shifts in genetic structure have been linked to population turnover and to fluctuations in population densities among years [56]. Prairie and meadow vole each display multi-annual population density cycles that are non-synchronous between species [50,57].

Vole captures in this study were not arrayed equally across years. Instead, a majority (47%) were collected in 2012, with 23% in 2011, and 30% in 2010. Population fluctuations were manifested across sites as well, with the smallest samples occurring for three sites in 2011, whereas two sites contained the fewest individuals in 2010 (Table 1). Lower densities will alter spatial structure at the metapopulation scale, it will be enhanced during periods of high densities [33], thus influencing dispersal tendencies and subsequently genetic structure.

#### Localized genetic structure

On a more local scale, genetic structure in voles can be attributed to the spatial clustering of kin [16] or reflect gender-specific differences in space use [56]. Prairie vole is monogamous, with strong pair bonds, and displays an elevated degree of philopatry and sociality [59]. Preferences for familiar peers are maintained in part by aggression toward unfamiliar individuals, as in mate partnerships [60]. Conversely, meadow vole is promiscuous and with males more likely to disperse [53]. Here, social tolerance is an important feature, as demonstrated by reduced aggression toward unfamiliar conspecifics [60].

At Prairie Ridge, clear genetic substructure was detected between two sampling plots separated by ~5km, and with restoration histories (intermediate *versus* established) that differed with regard to years since seeding [19]. The established plot (i.e., seeded >10 years ago) exhibited consistently elevated average relatedness when compared to the intermediate plot (seeded 3 years ago). This underscores that restoration age, as well as connectivity, may influence genetic structure. In fact, temporal stability of genetic composition in the root vole (*Microtus oeconomus*) was attributed in part to immigration of closely related individuals from nearby areas [54]. Thus, despite social structure (i.e., kin groups), gene flow may not significantly impact genetic composition of study population, despite large population fluctuations.

### Beyond restoration: Small mammals as disease vectors

Dispersal and population connectivity are aspects that also translate to a variety of ecological processes [61]. For example, they can help evaluate the porosity of the fragmented habitat to emerging infectious diseases (EIDs), and thus are key elements that can gauge availability of, and contact rates among, individuals of a host species [62].

Landscape genetic tools are often employed to characterize population dynamics, gene flow and movement pathways of disease hosts across heterogeneous landscapes [63], as well as to ascertain the concurrent spread and persistence of infectious agents associated with these movements [64]. For example, gene flow estimates in white-tailed deer (*Odocoileus virginianus*) corresponded to the rapid expansion of chronic wasting disease (CWD) in northern Illinois/ southern Wisconsin and served to identify those habitats with an elevated risk for infection [65].

In this regard, grassland rodents are recognized as important vectors and host reservoirs for EIDs, as evidenced by the contemporary spread of Hanta virus [66], as well as Lyme disease. Of interest in our study is the fact that Lyme disease does not impact the survival of the rodent [67], but merely utilizes them as a vector for the dissemination of the disease host (i.e., ticks). Importantly, Lyme disease is an EID of increasing concern due to its rapid geographic expansion promoted by climate change and has become a substantial threat to human health in midwestern North America [68]. It is one of the most common vector-borne illnesses in the United States (http://www.cdc.gov/lyme/stats/), with >30,000 cases reported annually. The expansion of Lyme disease is concomitant with the spread of the tick host (*I. scapularis*), as promoted by ongoing climate change [68]. Of importance is the capacity of *I. scapularis* to parasitizes multiple vectors that demonstrate both short- and long-distance dispersal [69]. Optimal habitat for the tick is prairie and young forest, and the prevalence of the causative agent for Lyme disease, *Borrelia burgdorferi* (*Bb*: the bacterium causing Lyme Disease), is 2x-greater in prairie habitat, with prairie vole the exclusive vector in those habitats [70].

Although we did not quantify the rate of *Bb* infection in our study individuals, it was previously demonstrated to be quite prevalent among prairie vole in our study area ([70]; see Fig. 1). In addition, the probability of transmission across restored grassland is quite high [71, 72], as suggested by the elevated genetic connectivity we found for prairie vole within plots (average width 3–7.5 km), and also among sites (at 48–246 km). This capacity for transmission, in turn, represents a largely unrecognized side effect that can develop in lockstep with the rehabilitation of agricultural lands. As such, it can represent an argument against a more positive interpretation of habitat restoration, with control measures for Lyme disease a primary concern [73]. Instead, our results, in tandem with non-significant *F_ST_* values, stand in contrast to the suggestion that well-separated habitat patches will potentially limit the spread of an EID.

## Conclusions

The restoration of agricultural plots to tall grass prairie is an ongoing process implemented in midwestern North America by federal and state resource agencies, but with success often being difficult to gauge. As a benchmark for successful management, we employed molecular ecology as a means to gauge restoration success by assaying for connectivity among vole populations found in rehabilitated Illinois prairie habitats. Successful habitat restoration may also elicit a potential downside in that it can increase the permeability of an environmental matrix (in this case, restored tallgrass prairie) to infiltration by vector-borne diseases.

## Author Contributions

MRD, RLS, MPM, and MED conceived and designed the experiments. WJBA, MAD, and MPM collected data. WJBA and MAD analyzed the data. WJBA, MAD, MRD, and MED wrote the manuscript.

## Acknowledgments

We thank P. Wolff, S. Beyer, W. Hill, S. McLaughlin, K. Barmann, C. Griffith, B. Neece, A. Ahlers, J. Andrews, G. Spyreas, and J. Larkin for fieldwork. Illinois Department of Natural Resources (IDNR) facilitated landowner access to SAFE sites. The U.S. Fish and Wildlife Service (USFWS) Federal Aid in Wildlife Restoration Program, as administered by the Illinois Department of Natural Resources (IDNR), provided funding for this project. Additional funding was provided by endowments through University of Arkansas/ Fayetteville to MRD (Bruker Professor of Life Sciences) and MED (21^st^ Century Chair in Global Change Biology). Opinions expressed herein represent those of the authors and do not reflect those of the Illinois Department of Natural Resources or the Illinois state government.

## Data accessibility

Microsatellite data are available on the Dryad Digital Repository (http://dx.doi.org/upon-acceptance).

## Author Contributions

**Conceptualization**: Marlis R. Douglas, Robert L. Schooley, Michael E. Douglas.

**Data curation**: Marlis R. Douglas, Whitney J. B. Anthonysamy, Mark A. Davis.

**Formal analysis**: Marlis R. Douglas, Whitney J. B. Anthonysamy, Mark A. Davis, Michael E. Douglas.

**Funding acquisition**: Marlis R. Douglas, Robert L. Schooley, Wade Louis, Michael E. Douglas.

**Investigation**: Marlis R. Douglas, Whitney J. B. Anthonysamy, Robert L. Schooley, Mark A. Davis, Robert L. Schooley, Wade Louis, Michael E. Douglas.

**Methodology**: Marlis R. Douglas, Whitney J. B. Anthonysamy, Mark A. Davis, Robert L. Schooley, Matthew P. Mulligan, Michael E. Douglas.

**Project administration**: Marlis R. Douglas, Robert L. Schooley, Wade Louis, Michael E. Douglas.

**Resources**: Whitney J. B. Anthonysamy, Mark A. Davis, Matthew P. Mulligan.

**Software**: Whitney J. B. Anthonysamy, Michael E. Douglas.

**Supervision**: Marlis R. Douglas, Robert L. Schooley, Wade Louis, Michael E. Douglas.

**Validation**: Marlis R. Douglas, Whitney J. B. Anthonysamy, Mark A. Davis, Michael E. Douglas.

**Writing – original draft**: Michael E. Douglas.

**Writing – review & editing**: Marlis R. Douglas, Whitney J. B. Anthonysamy, Mark A. Davis, Robert L. Schooley, Wade Louis, Matthew P. Mulligan, Michael E. Douglas.

## Supplemental Appendix I: Multiplex Panels

The 23 microsatellite DNA primers that successfully amplified loci in Prairie and Meadow Vole were combined into six multiplex panels for data generation. Included for each panel are: original primer name, repeat motif, citation, and forward and reverse primer sequences. Amplifications were conducted in 10-15 μl volume polymerase chain reactions (PCR) using 1X *Flexi*-buffer (Promega), 2.5-3.5 mM MgCl2, 0.25 mM dNTPs, 0.2 μg BSA, 1 unit *Go-taq* polymerase (Promega), 0.1 μM of each forward and reverse primer, and approximately 10-15 ng template DNA. Reactions were carried out under the following conditions: initial denaturation at 94°C for 3 min, followed by 15 cycles of 94°C for 45 s, 55°C annealing temperature for 45 s, and a 72°C extension for 30 s, followed by an additional 25 cycles of 95°C for 30 s, 55°C annealing temperature for 30 s, and a 72°C extension for 15 s, followed by a final extension at 72°C for 3 min.

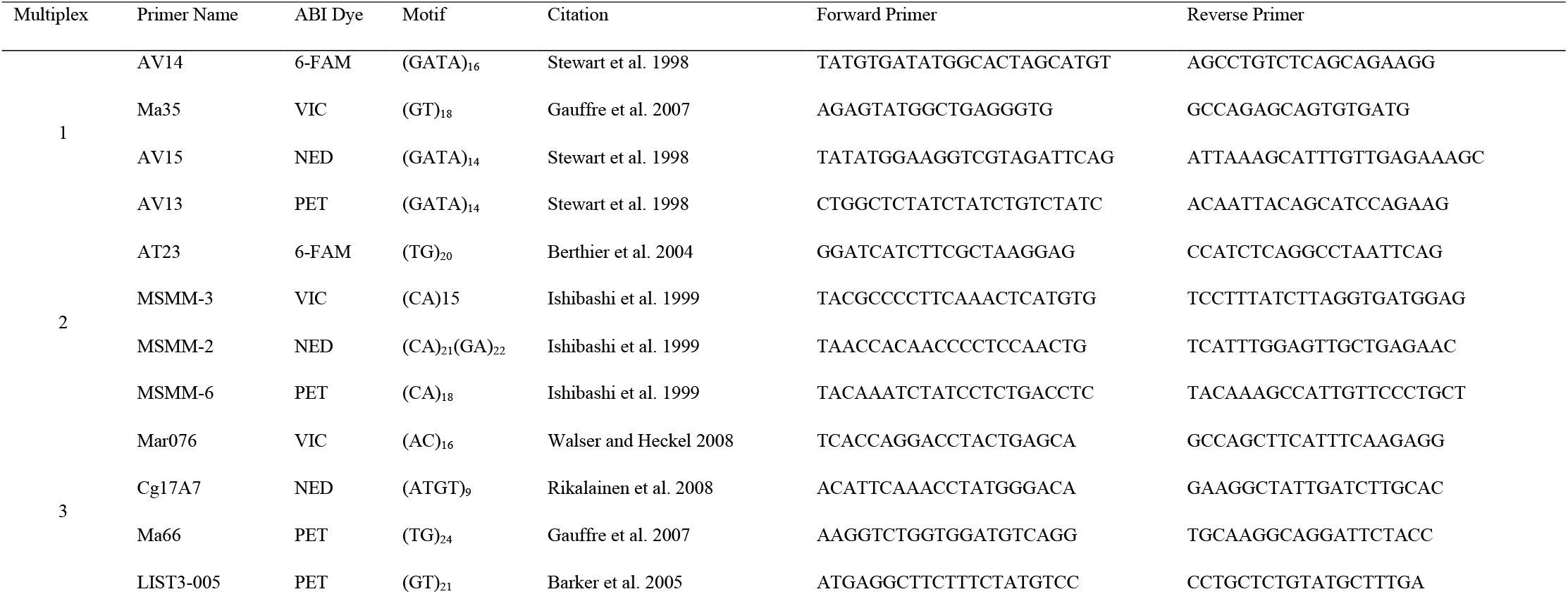
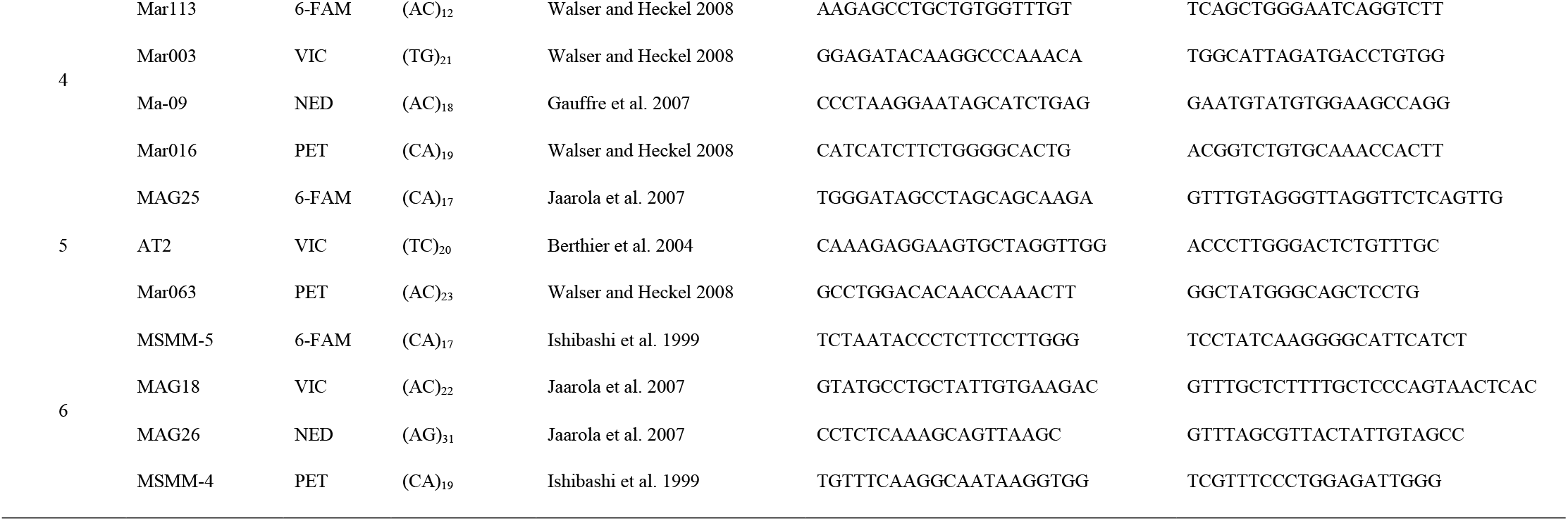

